# Freeze-dried porous collagen scaffolds for the repair of volumetric muscle loss injuries

**DOI:** 10.1101/2024.08.30.610194

**Authors:** Ivan M. Basurto, Ryann D. Boudreau, Geshani C. Bandara, Samir A. Muhammad, George J. Christ, Steven R. Caliari

## Abstract

Volumetric muscle loss (VML) injuries are characterized by the traumatic loss of skeletal muscle resulting in permanent damage to both tissue architecture and electrical excitability. To address this challenge, we previously developed a 3D aligned collagen-glycosaminoglycan (CG) scaffold platform that supported *in vitro* myotube alignment and maturation. In this work, we assessed the ability of CG scaffolds to facilitate functional muscle recovery in a rat tibialis anterior (TA) model of VML. Functional muscle recovery was assessed following implantation of either non-conductive CG or electrically conductive CG-polypyrrole (PPy) scaffolds at 4, 8, and 12 weeks post-injury by *in vivo* electrical stimulation of the peroneal nerve. After 12 weeks, scaffold-treated muscles produced maximum isometric torque that was significantly greater than non-treated tissues. Histological analysis further supported these reparative outcomes with evidence of regenerating muscle fibers at the material-tissue interface in scaffold-treated tissues that was not observed in non-repaired muscles. Scaffold-treated muscles possessed higher numbers of M1 and M2 macrophages at the injury while conductive CG-PPy scaffold-treated muscles showed significantly higher levels of neovascularization as indicated by the presence of pericytes and endothelial cells, suggesting a persistent wound repair response not observed in non-treated tissues. Finally, only tissues treated with non-conductive CG scaffolds displayed neurofilament staining similar to native muscle, further corroborating isometric contraction data. Together, these findings show that CG scaffolds can facilitate improved skeletal muscle function and endogenous cellular repair, highlighting their potential use as therapeutics for VML injuries.

## 1. Introduction

Volumetric muscle loss (VML) injuries result in an aberrant wound healing response that is characterized by chronic inflammation, persistent activation of myofibroblasts, excessive collagen deposition, and ultimately a loss of muscle mass and function^1–3^. These injuries are defined by the removal of a large volume of muscle tissue and the surrounding extracellular matrix (ECM) that is typically due to high energy trauma, such as blast injuries and vehicle accidents, surgical ablations, or myopathies^4^. The current clinical standard for treatment of VML injuries is autologous tissue transfer. However this process is time-intensive, expensive, and results in variable levels of functional success^5^. Furthermore, donor site morbidity as well as the heterogeneity between patients and injuries has made effective treatment an enduring challenge^6,7^. As a result, there is a clear need to design novel therapeutic approaches for VML injuries.

To address treatment limitations of VML, researchers have focused on the use of biomaterial systems to support endogenous repair mechanisms. These materials have varied widely to include natural and synthetic polymers augmented with growth factors, cells, and other biophysical cues. The considerable diversity of biomaterials used for skeletal muscle tissue engineering has been reviewed extensively elsewhere^8–12^. Careful design of biomaterial systems is critical to support improved regenerative outcomes because subtle changes to material properties can either facilitate regeneration or drive suboptimal pathological outcomes^13^. Two biophysical features that are characteristic of healthy skeletal muscle function are the high level of structural organization and the electrochemical excitability that enable efficient contraction, force generation, and locomotion. This is particularly relevant following VML where the ECM becomes disorganized, hindering cell-mediated repair^14^, and a prolonged loss of electrical stimuli results in continual muscle atrophy^15–17^.

In an effort to recreate these key biophysical features, we have previously developed 3D aligned collagen-glycosaminoglycan (CG) scaffolds for skeletal muscle tissue engineering^18,19^. CG scaffolds have a rich history of clinical use and were previously FDA-approved for skin regeneration^20,21^. Building on this platform, our previous work used a directional freezing process that resulted in aligned collagen struts reminiscent of healthy skeletal muscle ECM following lyophilization^22^. To recapitulate the electrical properties of skeletal muscle, we also demonstrated the ability to synthesize and incorporate electrically-conductive polypyrrole (PPy) particles into our CG suspension prior to freeze-drying. The inclusion of PPy resulted in a five-fold increase in material conductivity that did not detrimentally impact myoblast metabolic activity. Moreover, our PPy-doped scaffolds supported 3D cytoskeletal alignment along the scaffold’s backbone and promoted enhanced myoblast maturation measured by myosin heavy chain (MHC) expression. As a logical extension of our prior work, we aim to explore how 3D aligned collagen scaffolds, in both non-conductive and electrically-conductive forms, influence skeletal muscle repair in an *in vivo* rat VML injury model.

## 2. Materials and Methods

### 2.1 Polypyrrole (PPy) synthesis

PPy nanoparticles were synthesized as previously described^18^. Briefly, 2 g of pyrrole monomer was reacted with 72 mmol FeCl_3_ using vigorous mixing for 24 h under ambient conditions. The resulting black precipitate was washed repeatedly with water and filtered using vacuum filtration. The PPy powder was then dried overnight and passed through a 325-mesh (45 μm) screen. Fourier-transform infrared (FTIR) spectroscopy was then used to confirm the chemical structure of the PPy particles.

### 2.2 Scaffold fabrication

Non-conductive collagen-glycosaminoglycan (CG) scaffolds were fabricated by homogenizing a suspension of 1.5 wt% microfibrillar type I collagen from bovine Achilles tendon and 0.133 wt% chondroitin sulfate derived from shark cartilage in 0.05 M acetic acid. The suspension was prepared in a recirculating chiller maintained at 4°C to prevent collagen denaturation. Conductive PPy-doped (CG-PPy) scaffolds were created by mixing PPy nanoparticles (0.5 wt%) into the collagen/chondroitin sulfate suspension via vortexing. All scaffolds were fabricated via directional lyophilization using a thermally mismatched mold^18,23^ in a VirTis Genesis pilot scale freeze-dryer. Following lyophilization scaffolds were dehydrothermally crosslinked at 105°C for 24 h.

### 2.3 Scaffold hydration and crosslinking

The resulting scaffold cylinders (∼ 15 mm height, ∼ 6 mm diameter) were hydrated in 70% ethanol for 30 min before being transferred to PBS. The scaffolds were then chemically crosslinked using 1-ethyl-3-(−3-dimethylaminopropyl) carbodiimide hydrochloride (EDC) and N-hydroxysulfosuccinimide (NHS) at a molar ratio of 5:2:1 EDC:NHS:COOH where COOH is the carboxylic acid content of the collagen. EDC/NHS-mediated crosslinking facilitates the covalent reaction of collagen primary amines with carboxylic acids to improve scaffold mechanical integrity^24^. CG scaffolds were incubated in sterile-filtered EDC/NHS solution for 50 min under moderate shaking before being washed twice with PBS. The scaffolds were then transferred to 70% ethanol to sterilize overnight before being washed repeatedly with sterile PBS. All scaffolds were stored at 4°C in sterile PBS until the time of surgery.

### 2.4 Animal care

This study was conducted in compliance with the Animal Welfare Act, the Implementing Animal Welfare Regulations, and in accordance with the principles of the Guide for the Care and Use of Laboratory Animals. The University of Virginia Animal Care and Use Committee approved all animal procedures. A total of 24 male Lewis rats (Charles River Laboratories) age-matched to 11 weeks weighing 312.7 ± 24.9 g were purchased and individually housed in a vivarium accredited by the American Association for the Accreditation of Laboratory Animal Care and provided with food and water *ad libitum*.

### 2.5 Surgical procedures

The VML injury model was created using a previously established tibialis anterior (TA) injury model^25,26^. Rats were randomly designated to three different experimental groups: no repair (NR; *n* = 8), non-conductive CG scaffolds (CG; *n* = 8), and conductive PPy-doped CG scaffolds (CG-PPy; *n* = 8). To create the surgical defect, rats were anesthetized via isoflurane and the surgical site was aseptically prepared by repeated washes with alcohol and iodine. A longitudinal skin incision was made on the anterior side of the lower left leg to expose the anterior crura muscles. The skin was then separated from the underlying fascia using surgical scissors. The fascia was then separated to expose the underlying musculature. Next, the extensor digitorum longus (EDL) and extensor hallucis longus (EHL) muscles was surgically ablated to avoid compensatory hypertrophy of synergistic muscles involved in dorsiflexion during later *in vivo* functional assessment^25^. The VML defect size was approximated from a validated linear regression to estimate TA muscle mass from rat body weight^25,26^. The injury was created by surgically resecting approximately 20% of the TA muscle weight from the middle third of the TA muscle. Following creation of the injury, scaffolds were sutured into the defect using 6-0 Vicryl. The fascia was then sutured back in place using 6-0 Vicryl sutures, and the skin was closed with 5-0 Prolene using interrupted sutures. Skin glue was applied over top of the Prolene sutures to avoid reopening of the injury. Following surgery animal health was monitored daily and skin sutures were removed after 14 days.

### 2.6 In vivo functional testing

Isometric force testing was performed on animals at 1 week prior to surgery and at 4, 8, and 12 weeks post-surgery to quantify the functional deficit created by the surgical defect and track recovery over time. *In vivo* force testing was conducted using an established protocol in which isometric torque is produced as a function of stimulation frequency (1-200 Hz)^27^. Rats were anesthetized via isoflurane and the left hindlimb was aseptically prepared by repeated washes with alcohol and iodine. The foot was secured against a force transducer foot plate, ensuring that the heel was flush against the bottom of the plate. The knee joint was then stabilized and positioned so the foot and tibia were at a 90° angle. Force testing was performed by stimulation of the peroneal nerve with platinum needle electrodes. Electrodes were placed in the posterior compartment of the lower leg along either side of the peroneal nerve and muscle length was adjusted until maximal twitch force was produced. The contractile force of the anterior crura muscles was then assessed by measuring peak isometric tetanic force production as dorsiflexion occurred. Torques at all timepoints were normalized to the body weight of the animal at the time of stimulation.

### 2.7 Histology analysis

Following functional testing all retrieved muscles were photographed before being frozen for further processing. The TA muscle was cut in half cross-sectionally, and the proximal portion was embedded in OCT before being flash frozen in liquid nitrogen-cooled isopentane. Distal muscle sections were cut in half longitudinally and embedded in OCT for cryosectioning. Distal sections were used for analysis of muscle innervation while proximal tissue sections were used to quantify muscle fiber cross sectional area (FCSA), macrophage infiltration, and neovascularization. All muscle samples were then cryosectioned (8 μm-thick slices) and placed on glass slides. Hematoxylin and Eosin (H&E) stains were administered using conventional techniques to analyze tissue morphology, muscle repair, and fibrotic response for at least three muscles per experimental group. Stained muscle sections were then visualized using a Leica Thunder imaging system at 10x magnification.

### 2.8 Immunohistochemical (IHC) analysis

Unstained OCT-embedded slides were first subjected to an antigen retrieval protocol (H-3301; Vector Laboratories) in preparation for antibody staining. To reduce autofluorescence of muscle tissue, samples were treated with 0.3% Sudan black solution for 10 min prior to blocking for 2 h (Dako Blocking Solution X0909; Agilent Technologies, Santa Clara, CA). Immunohistochemical staining was then performed using antibodies to detect laminin (dilution 1:200, ab11575; Abcam), CD68 (1:100, MCA341R; Bio-Rad) and CD163 (1:400, ab182422; Abcam), CD31 (1:250; Novus Biologicals NB100-2284) and α-smooth muscle actin (αSMA, conjugated to 488 fluorochrome, F3777, 1:250; Sigma Aldrich), or neurofilament 200 (NF200, anti-chicken, 1:1000; EnCor CPCA-NF-H) overnight at 4°C. Next, samples were incubated for 2 h at ambient temperature with one or more of the following secondary antibodies: Alexa Fluor 647 Fab 2 fragment goat anti-rabbit (1:500), Alexa Fluor 488 goat anti-chicken (1:600), Alexa Fluor 488 goat anti-mouse (1:400), and/or Alexa Fluor 488 goat anti-rabbit (1:400). Finally, slides were stained with DAPI (1:1000) before being stored in a light-protected environment at -20 °C until imaging.

All samples were imaged on a Leica inverted confocal microscope using a 10x objective across the entire TA muscle section (*n* = 3 muscles for each group). For quantification of macrophage infiltration and polarization, image analysis was conducted at the injury site, defined as a 4 mm x 2 mm region at the VML injury. A custom MATLAB code was then used to quantify the number of M1 (CD68^+^/CD163^-^) and M2 (CD163^+^) macrophages throughout this region. CD31 and αSMA-labeled structures were quantified using CellProfiler. For neurofilament-stained images, NF200^+^ structures were quantified across the whole longitudinal muscle sections using ImageJ.

### 2.9 Muscle fiber minimum Feret diameter quantification

Muscle sections stained with laminin were used to visualize the ECM and quantify minimum Feret diameter (the measure of an object size along its minimum axis), fiber cross-sectional area (FCSA), and total fiber count. Laminin-stained muscle sections were visualized using a Leica inverted confocal microscope DMi8 with a 5x objective. Image post-processing was conducted using ImageJ. Muscle fiber quantification was conducted using a publicly available semi-automatic muscle analysis using segmentation of histology (SMASH) software^28^. Briefly, individual muscle fibers were outlined using SMASH’s built-in segmentation algorithm based on laminin staining. The segmentation output was visually inspected and incomplete or incorrect segmentation was manually adjusted. Following correct segmentation, the number of muscle fibers and FCSA were calculated by the software across the whole TA muscle section. To gain a more thorough understanding of the quality of muscle repair, SMASH analysis was repeated at the VML injury site (4 mm x 2 mm region at the VML injury) to quantify minimum Feret diameter and FCSA across experimental groups. The analysis was completed in triplicate for each experimental group.

### 2.10 Statistical analysis

Data are presented as means and their standard deviations (SDs) unless otherwise indicated. Histological, immunohistochemical, and muscle fiber quantification was conducted for *n* = 3 muscles per group. Data normality distribution was evaluated using the D’Agostino and Pearson test. Kruskal-Wallis with Dunn’s multiple comparisons tests were performed for data that were not normally distributed. Functional data were statistically analyzed using a paired two-way analysis of variance (ANOVA) while immunohistochemical staining was evaluated using a one-way ANOVA. Upon finding any statistically significant differences via ANOVA, post-hoc multiple comparison tests of parameters of interest were performed using Tukey’s HSD or Dunnett’s multiple comparisons test. These statistical analyses were conducted using GraphPad Prism 9.0. *P* values < 0.05 were considered statistically significant.

## 3. Results

### 3.1 Creation of rat TA VML injury and scaffold delivery

CG scaffolds were surgically delivered into an established rat tibialis anterior (TA) model of volumetric muscle loss (VML) injury. The weight of excised muscle was statistically similar across experimental groups, indicating reproducible creation of VML injuries. Three experimental groups were investigated: No repair (NR: 105.3 ± 11.6 mg, *n* = 8) and implantation of either non-conductive CG scaffolds (CG: 114.6 ± 14.2 mg, *n* = 8) or conductive PPy-doped CG scaffolds (PPy: 116.0 ± 7.8 mg, *n* = 8) (**Figure 1**). Following surgical creation of the defect, scaffolds were cut to the dimensions of the injury and sutured into place. No deaths occurred following the procedure and all animals recovered with no visual signs of infection or additional treatment. Moreover, animal body weights were statistically similar at the time of surgery and underwent comparable increases over the course of the study.

**Figure 1:**
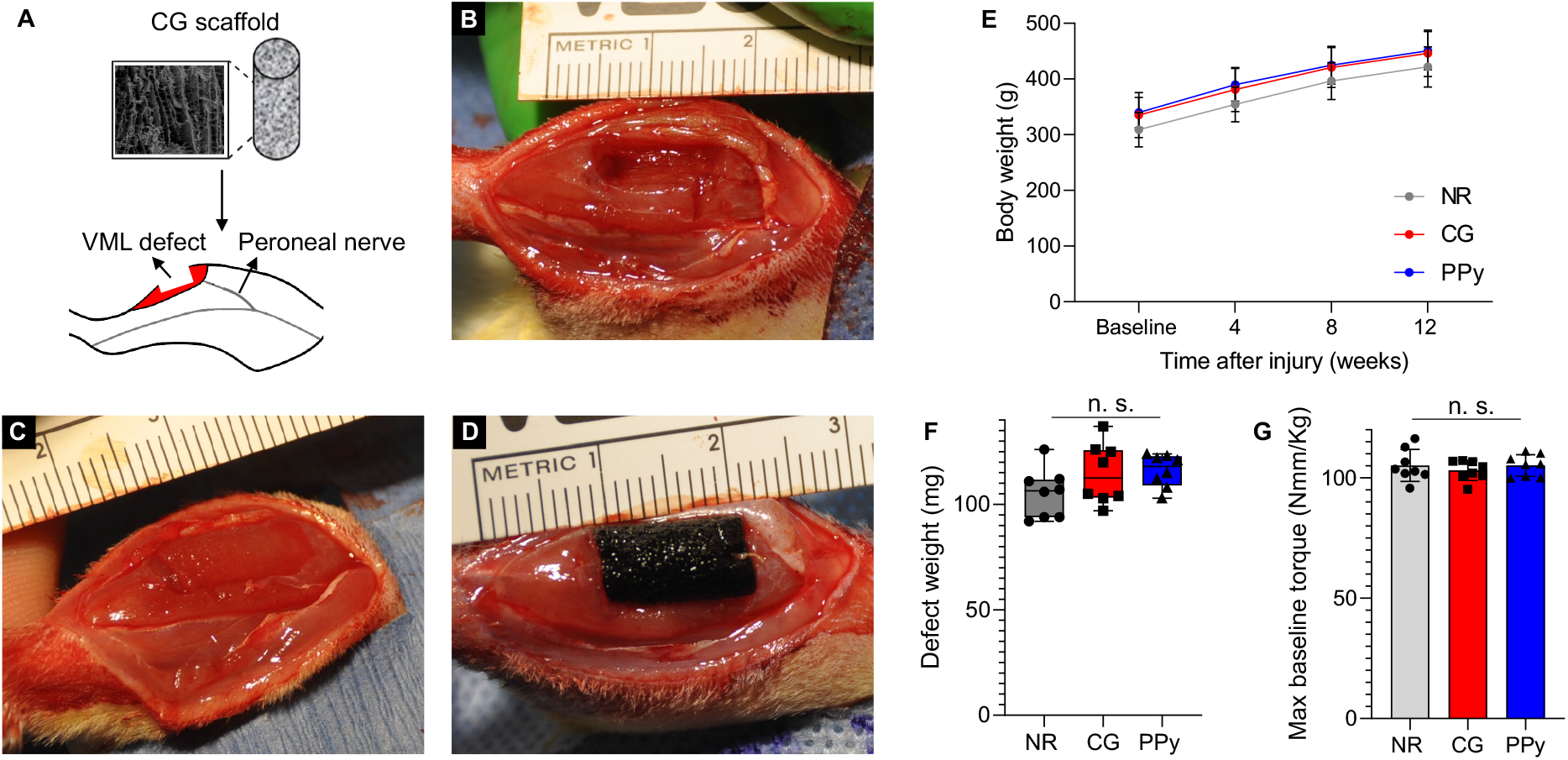
CG scaffolds were surgically implanted into a tibialis anterior (TA) model of VML injury. (A) Schematic representation of scaffold delivery into TA VML injury model. Representative images of (B) no repair, (C) non-conductive CG scaffold and (D) PPy-doped CG scaffold-treated muscles. (E) Animal body weight 1 week prior to surgery and at 4, 8, and 12 weeks post VML injury, corresponding to functional testing time points. (F) Weight of defects created for no repair, non-conductive CG scaffold, and conductive PPy-doped CG scaffold (NR, CG, and PPy, respectively) experimental groups. (G) Maximum baseline torque generation normalized to animal body weight pre-injury. Data presented as Mean +/-SD (panels E and G) while panel F data are presented as box plots of interquartile range (line: median) with whiskers showing minimum and maximum values. n.s.: no statistically significant differences. *n* = 8 animals per experimental group.

### 3.2 Evaluation of TA functional recovery post-VML injury

Functional muscle contraction was assessed prior to surgery and at 4, 8, and 12 weeks post-surgery by *in vivo* stimulation of the TA muscle. Briefly, the animal’s left hindlimb was affixed to a force transducer and electrical probes were inserted along the peroneal nerve. Isometric contractile force in response to direct muscle electrical stimulation was recorded over a range of frequencies (1-200 Hz). Force production was reported as torque normalized to body weight at each time point (N-mm kg^-1^ of body weight) to account for increases in force production due to animal growth. There were no statistical differences in maximum baseline isometric torque between experimental groups prior to surgery. Additionally, baseline values reflect our previously published data using similar methods^29–31^. At 4 weeks post-VML there were no statistical differences in torque production amongst the treatment groups (**Figure 2**). NR and scaffold-treated muscles produced a maximum isometric torque that was roughly 50% of baseline values (NR: 49.5 ± 5.7 N-mm kg^-1^, CG: 54.8 ± 3.8 N-mm kg^-1^, PPy: 55.3 ± 7.8 N-mm kg^-1^) and was similar to previously published results^29–31^. At 8 weeks following injury, non-conductive CG scaffold-treated animals showed significantly improved isometric contraction at higher stimulation frequencies (60-200 Hz) compared to NR animals. When peak isometric torque was normalized to baseline values, CG scaffold-treated animals produced significantly greater force (62.7 ± 4.9%) than NR animals (52.3 ± 5.9%). A similar trend was observed in conductive CG-PPy-treated animals, although these results were not statistically significant. Finally, at 12 weeks post-injury both conductive CG-PPy and non-conductive CG scaffold-treated animals produced significantly greater isometric torque at higher stimulation frequencies (100-200 Hz, 60-200 Hz respectively) compared to non-treated animals. This trend was conserved when peak isometric torque was normalized to baseline values (NR: 51.6 ± 6.4%, CG: 64.5 ± 8.2%; PPy: 62.0 ± 5.7%) indicating superior functional muscle recovery by the addition of collagen scaffolds compared to non-treated animals.

**Figure 2:**
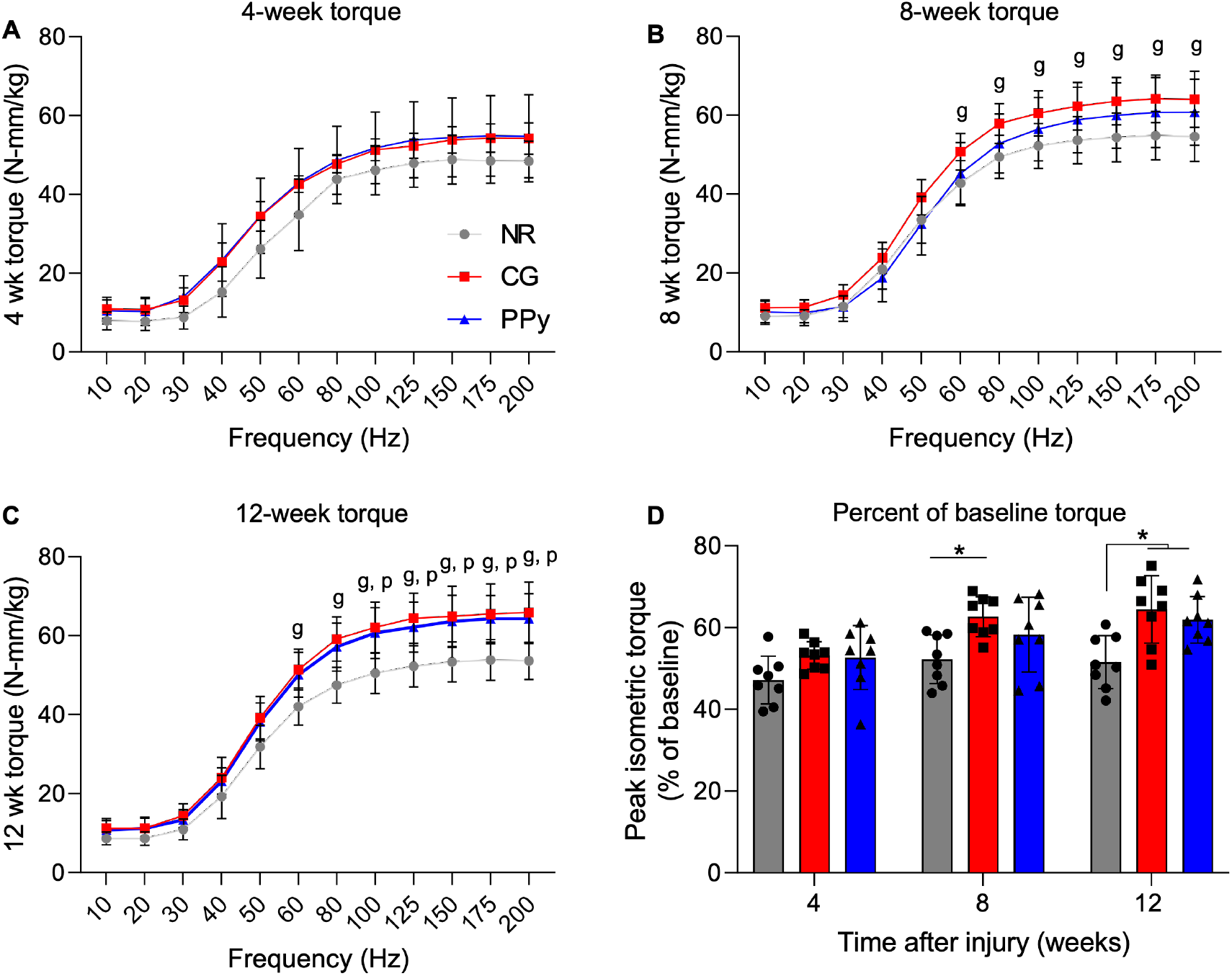
Both conductive and non-conductive scaffolds supported improved functional muscle recovery at 12 weeks post-VML. (A) Torque vs. frequency curves at 4 weeks post-injury show a roughly 50% drop in muscle force from baseline values. (B) At 8 weeks post-VML, non-conductive scaffold-treated muscles showed significantly increased torque production at higher stimulation frequencies. (C) At 12 weeks post-injury, conductive CG-PPy and non-conductive CG scaffold-treated tissues produced significantly more force in response to higher stimulation frequencies. (D) Peak isometric contraction normalized to baseline values showed that both CG-PPy and CG scaffolds facilitated increased functional recovery compared to non-treated tissues. Data presented as Mean +/-SD. Statistically significant differences compared to the no repair group are denoted by *g* (CG) and *p* (PPy). * *P* < 0.05. *n* = 8 animals per experimental group.

### 3.3 Gross tissue morphology

Following functional testing at 12 weeks post-VML, the TA of both injury and contralateral limbs were surgically explanted for morphological and histological analysis. Non-repaired muscles possessed a visible layer of fibrotic tissue and fatty deposition over the VML injury area (**Figure 3**). Additionally, visual inspection of non-treated muscles showed a convex surface at the defect compared to native contralateral control tissues. In contrast, non-conductive CG scaffold-treated muscles contained lower levels of ECM deposition, although some fibrotic tissue remained. Importantly, a clear discontinuity between the muscle tissue and implanted scaffold was not observed, indicating some degradation and integration of the CG scaffold. Residual PPy particles were clearly observed at the VML injury site in conductive scaffold-treated muscles 12 weeks post-injury.

**Figure 3:**
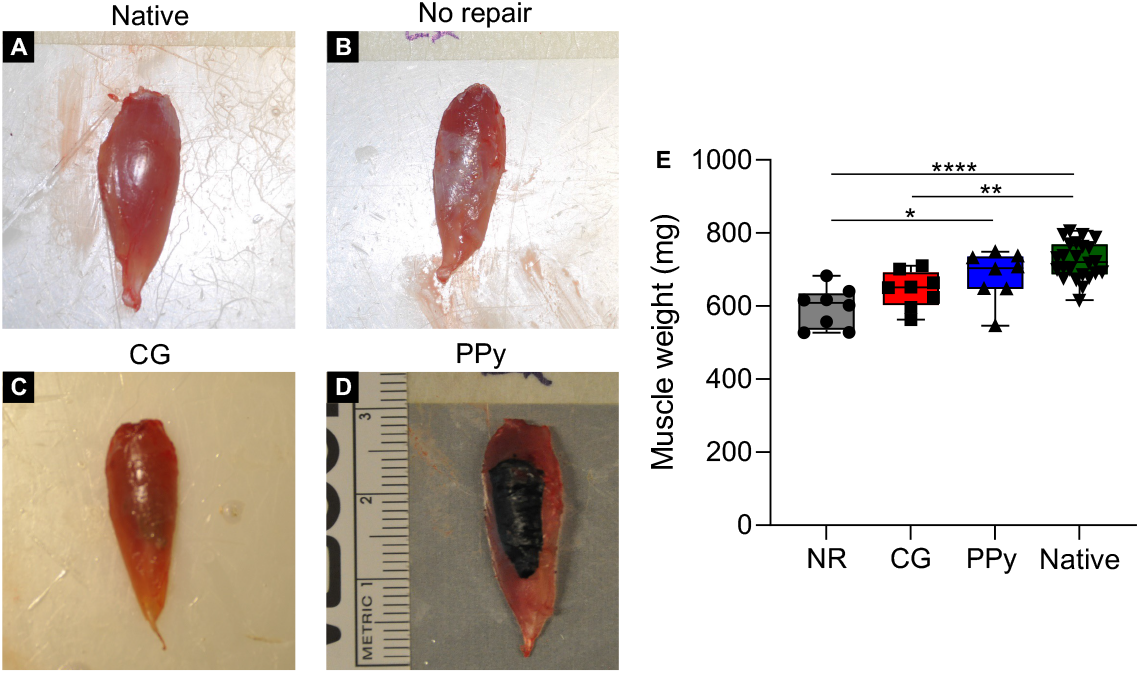
Gross tissue morphology at 12 weeks post-VML indicates a disparate biological response to therapeutic intervention. (A) Representative image of uninjured native tissue. (B) Non-repaired muscles possessed a layer of fibrotic and fatty tissue that surrounded the VML injury area. (C) CG scaffold-treated muscles displayed lower levels of fibrosis, but some ECM deposition was observed. (D) PPy particles remained localized to the VML defect, although the collagen scaffold appeared degraded. (E) Muscle weight at the time of explant shows that NR and CG scaffold-treated tissues were significantly lighter than native muscle. Data presented as box plots of interquartile range (line: median) with whiskers showing minimum and maximum values. * *P* < 0.05, ** *P* < 0.01, **** *P* < 0.0001. *n* = 8 muscles per experimental group and *n* = 24 for the native contralateral control.

While the particles remained, the collagen scaffold backbone appeared to be fully degraded as in CG scaffold-treated muscles. Evaluation of explanted TA muscle mass revealed that non-treated and CG scaffold-treated tissues (NR: 596.3 ± 55.2 mg, CG: 645.0 ± 49.8 mg), but not CG-PPy-treated tissues (PPy: 683.4 ± 67.5 mg), were significantly lighter than native uninjured muscles (Native: 723.9 ± 50.3 mg, *n* = 24), suggesting a persistent loss of muscle volume. CG-PPy scaffold-treated muscles were significantly heavier than non-treated tissues, although this is likely due to the continued presence of residual PPy.

### 3.4 Histological analysis

After functional testing, explanted muscles were frozen and embedded for histological and immunohistochemical analysis. Muscles explanted from three animals per experimental group were transversely sectioned and underwent H&E staining to analyze muscle morphology and fibrotic response following VML injury. Native tissue sections showed an organized network of muscle fibers with nuclei along the periphery of the myofibers characteristic of healthy muscle (**Figure 4**). As observed from the gross tissue morphology, the VML defect area was easily identified in NR muscles as a clear concave absence of tissue volume. Moreover, there was a distinct lack of myofibers with centrally-located nuclei, a hallmark of regenerating muscle, suggesting limited muscle repair. In contrast, scaffold-treated muscles showed myofibers with centrally-located nuclei along the material-tissue interface, indicating myogenesis. Closer inspection of CG scaffold-treated muscles revealed a thin layer of connective tissue at the site of material implantation, as indicated by pink fibers, dispersed with cells. Together, these data indicate that while non-conductive scaffolds supported some level of myogenesis, the scaffold alone could not completely halt the fibrotic response characteristic of VML. Similarly, CG-PPy-treated muscles possessed evidence of regenerating myofibers with centrally located nuclei and a layer of connective tissue at the implantation site. H&E staining also showed that cells infiltrated within the residual PPy particles, suggesting a persistent cellular response and tissue remodeling.

**Figure 4:**
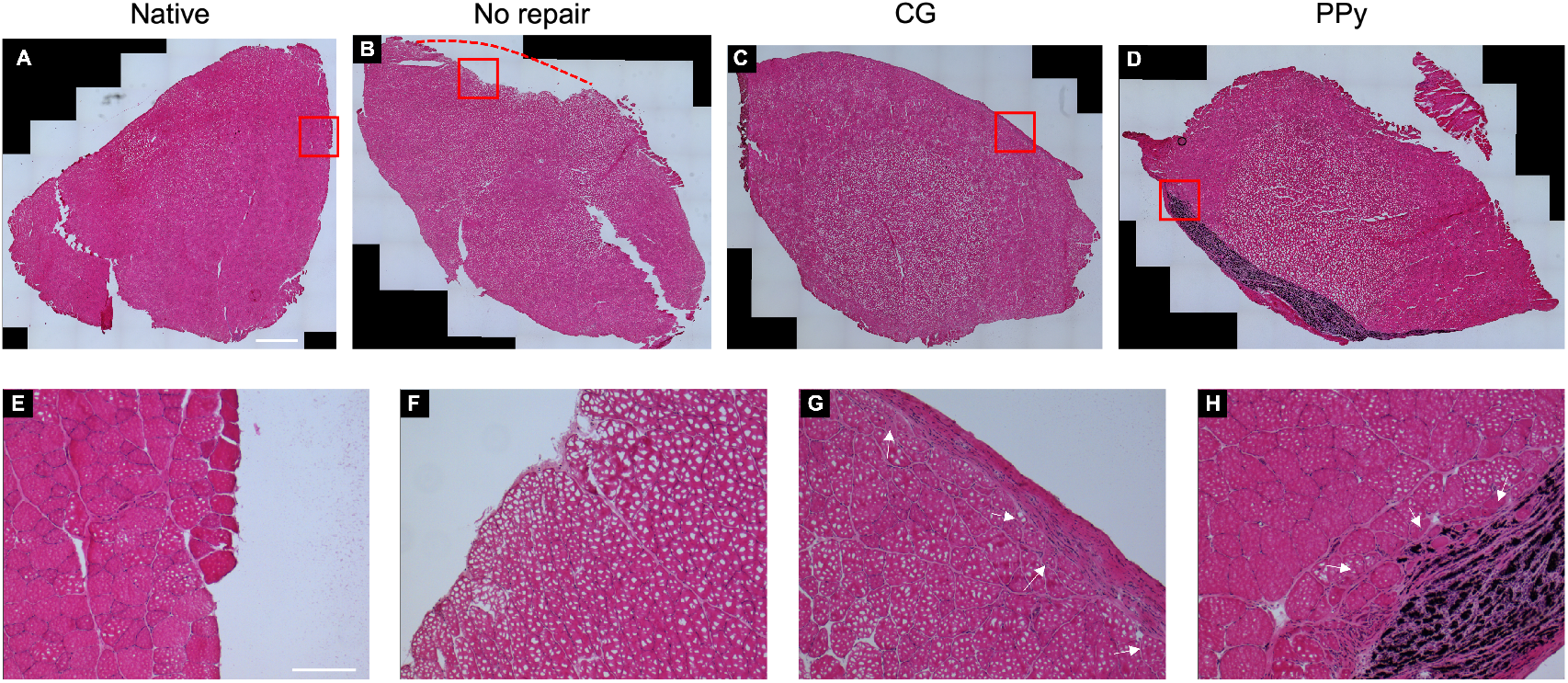
Histological analysis of TA muscles at 12 weeks post-injury. (A) H&E images of TA muscle cross sections for uninjured native tissue, (B) no repair, (C) CG scaffold, and (D) PPy-doped scaffold experimental groups at 12 weeks post-VML. *Dashed red line* indicates the concave VML defect area that remained due to limited tissue regeneration in the no repair group. *Red squares* denote regions of magnified images (E-H). (E) Magnified views of native tissue, (F) no repair, (G) CG scaffold, and (H) CG-PPy scaffold experimental groups. *Arrows* indicate myofibers with centrally located nuclei. Scale bars: 1 mm (top), 200 μm (bottom).

### 3.5 Characterization of regenerating muscle fiber cross-sectional area

To characterize the extent of muscle repair more thoroughly, tissue sections were stained for the ECM protein laminin to visualize individual muscle fibers. Semi-automatic muscle analysis using segmentation of histology (SMASH) software was then used to quantify the number of muscle fibers, fiber cross-sectional area (FCSA), and minimum Feret diameter^30^. Analysis of the total number of myofibers showed a reduction in fiber counts for all experimental groups compared to native muscles, although these results were not statistically significant (NR: 7858 ± 1420, CG: 8077 ± 1235, PPy: 7373 ± 434, Native: 10614 ± 852; **Figure S1**. The analysis was then repeated for muscle fibers at the location of VML injury, a 4 mm x 2 mm region at the tissue-injury interface, to evaluate regenerating muscle myofibers. Similarly, the number of myofibers at the VML injury site was not statistically different across experimental groups (**Figure 5**). Assessment of median FCSA and minimum Feret diameter (**Figure S2**) revealed that myofiber size was significantly reduced in non-treated muscles (1511, 607, 3031 μm^2^; reported as median, 1^st^, and 3^rd^ quartiles) compared to native tissue (3397, 1896, 5247 μm^2^), indicating muscle atrophy. In contrast, CG and CG-PPy scaffold-treated muscles possessed statistically similar median FCSA (CG: 2080, 778, 3847 μm^2^; PPy: 1617, 603, 4137 μm^2^) compared to uninjured muscles, further corroborating the improved functional outcomes observed. Characterization of FCSA at the VML site also revealed that muscle fibers were generally smaller in experimental tissues compared to native uninjured tissues. This trend is apparent by the increased frequency of smaller diameter fibers and decreased presence of larger muscle fibers compared to fibers in native tissues.

**Figure 5:**
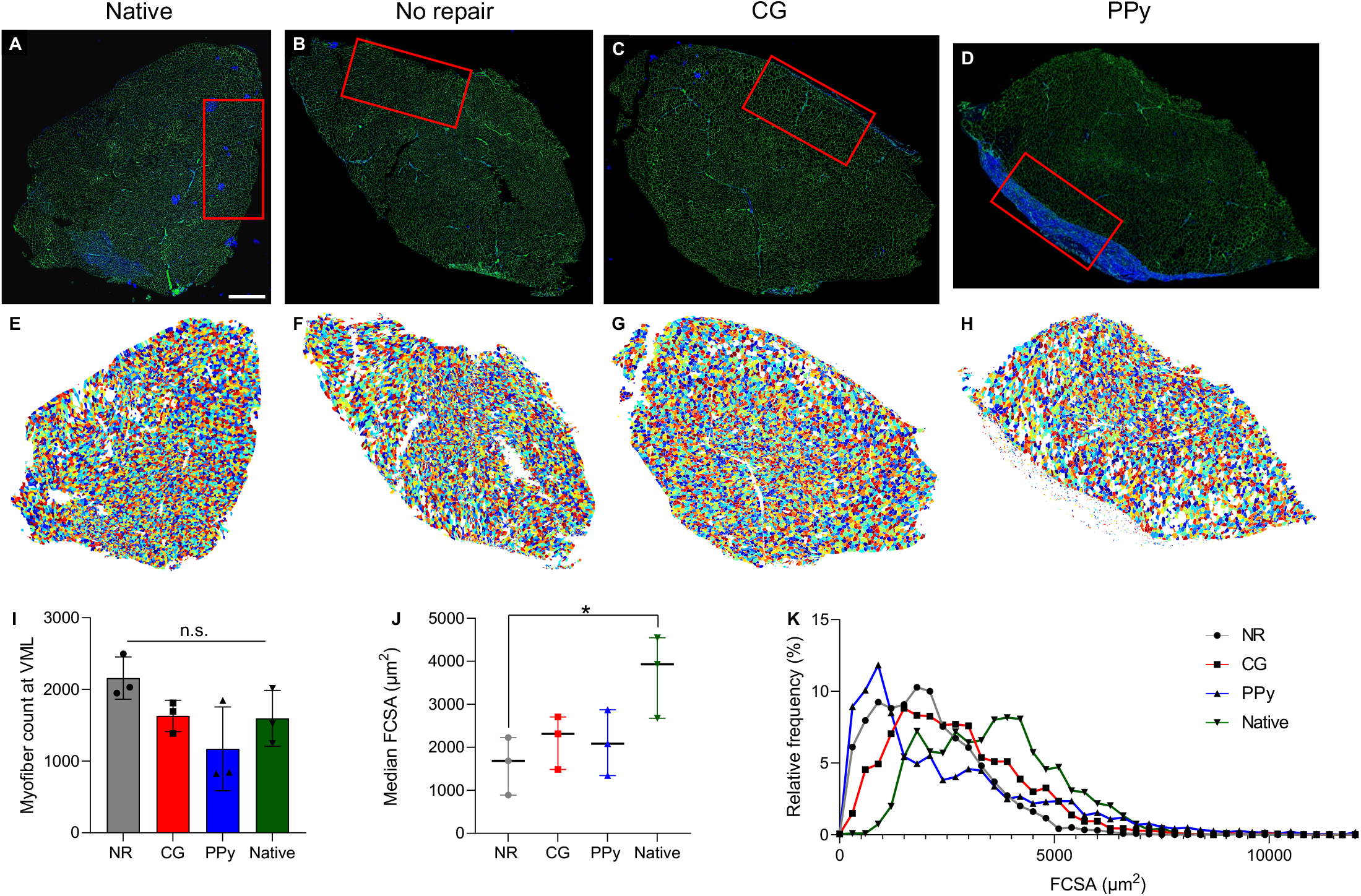
Myofiber cross-sectional area is significantly reduced in non-treated muscle tissues. (A) Representative laminin-stained sections of TA muscles for native uninjured control, (B) no repair, (C) non-conductive CG scaffolds, and (D) conductive CG-PPy scaffold-treated muscles at 12 weeks post-VML. *Red rectangles* denote region of VML injury used for FCSA analysis. (E–H) Colorized outputs show individual myofibers from SMASH software corresponding to (A–D), respectively. (I) Muscle fiber count at the VML injury was not statistically different from native muscle across experimental groups. (J) Median fiber cross-sectional area (FCSA) was significantly reduced in NR muscles. (K) FCSA relative frequency curves show a leftward shift toward smaller muscle fibers compared to uninjured muscle regardless of treatment type. Data presented as Mean +/-SD while panel (J) data are presented as the median (line: median) with interquartile range (whiskers). n.s.: no statistically significant differences. *n* = 3 muscles per experimental group. Scale bar: 1 mm.

### 3.6 Evaluation of long-term macrophage polarization

For minor muscle injuries, a pro-inflammatory immune response is activated within the first few days following injury that is gradually attenuated to allow for myoblast proliferation and differentiation into mature skeletal muscle. However, in the context of VML, a persistent immune response mediated by inflammatory cells such as macrophages results in excessive myofibroblast activation, minimal myogenesis, and limited functional repair^3^. To more probe how implanted scaffolds impacted the immune response following VML injury, macrophage infiltration and polarization was evaluated. Macrophage presence within the VML injury was characterized using CD68, a pan macrophage marker, and CD163, a marker of alternatively-activated M2 macrophages (**Figure 6**)^32^. The number of a classically-activated M1 pro-inflammatory macrophages (CD68^+^/CD163^-^) remained elevated in CG and CG-PPy scaffold-treated muscles compared to contralateral control muscles, although these results were not statistically significant. The continued presence of M1 macrophages may indicate a persistent inflammatory response^33,34^. Interestingly, the quantity of CD68^+^/CD163^-^ macrophages within non-repaired muscles was most similar to native muscle tissues, suggesting that other inflammatory cell types may be the primary drivers of chronic inflammation characteristic of VML. The expression of CD163, indicative of alternatively-activated M2 macrophages, was not statistically different across experimental groups at 12 weeks post-injury, although their numbers remained elevated in scaffold-treated tissues^35–37^.

**Figure 6:**
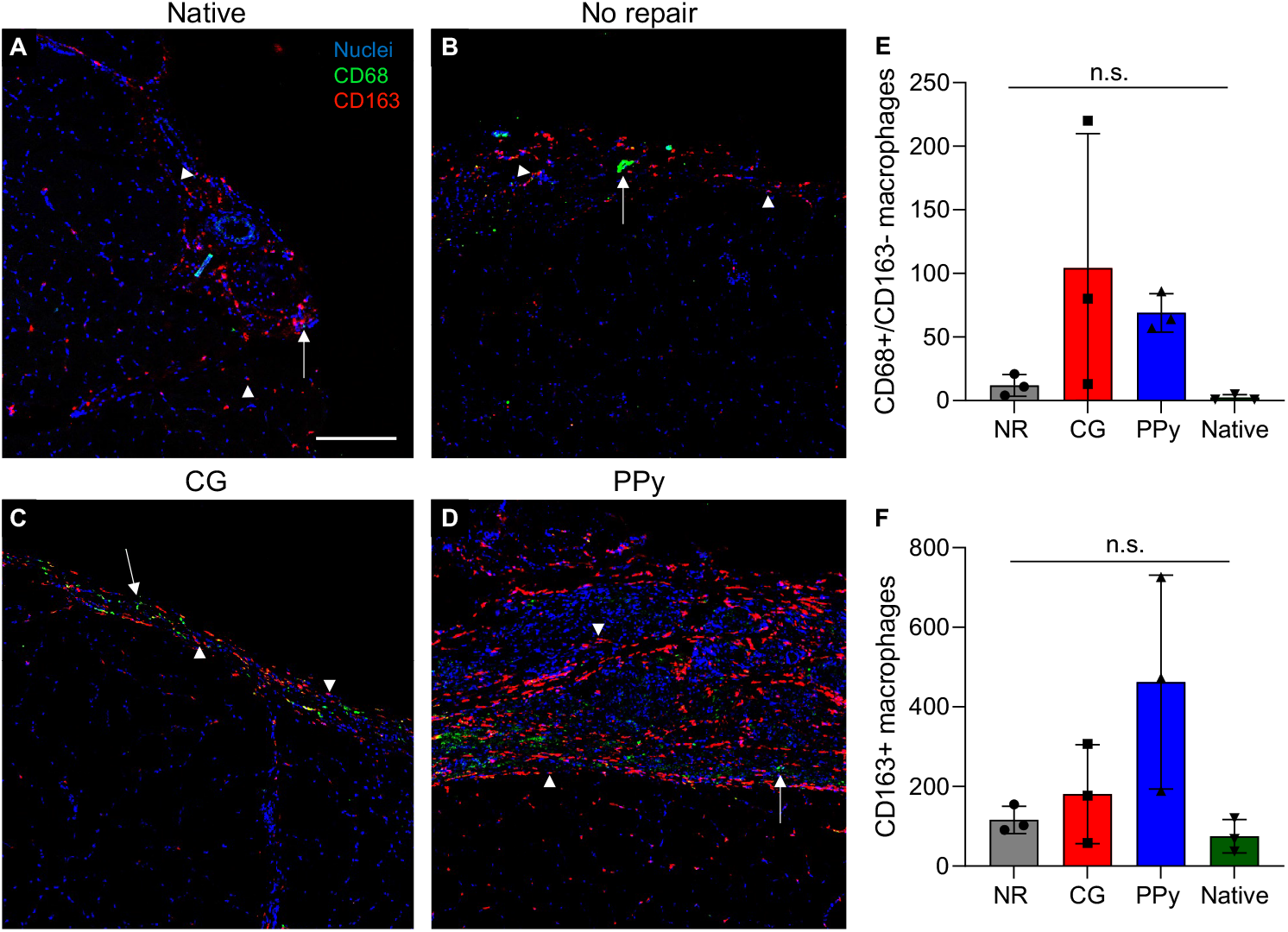
Scaffold-treated muscles show elevated macrophage infiltration at VML injury compared to native muscle. (A-D) Representative images of macrophage infiltration at 12 weeks post-VML. Images were taken at the site of VML injury and stained for CD68 (*green*, arrow) and CD163 (*red*, arrowhead). (E) CD68^+^/CD163^-^ M1 macrophages remained elevated in CG and CG-PPy scaffold-treated muscles, indicating a prolonged wound healing response, although these results were not statistically significant. (F) Quantification of CD163^+^ cells indicative of M2 macrophages showed no statistically significant differences across experimental groups. Data presented as Mean +/-SD. n.s.: no statistically significant differences. *n* = 3 muscles per experimental group. Scale bar: 200 μm.

### 3.7 Assessment of vascularization

Vascularization of muscle samples 12 weeks post-VML was evaluated using IHC staining. The presence of endothelial cells was examined through CD31 staining, while larger vessels were identified by simultaneous positive detection of CD31 and alpha smooth muscle actin (αSMA), which serves as a marker for pericytes. Comparing levels of CD31 and αSMA in the experimental groups to those in native muscle tissue, CG-PPy scaffold-treated muscles exhibited significantly higher levels of both CD31^+^ cells and αSMA structures (**Figure 7**).

**Figure 7:**
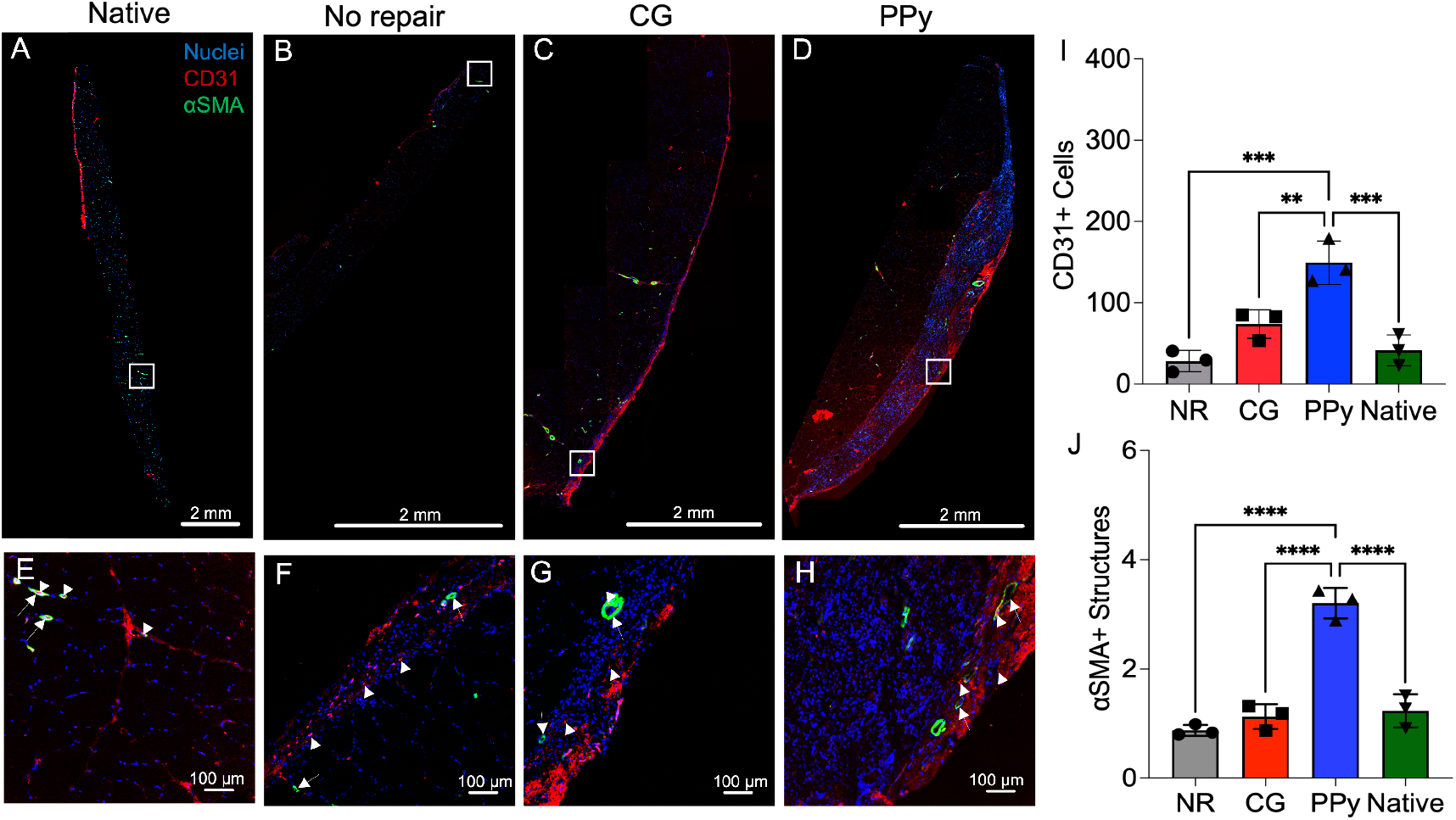
Conductive scaffolds support higher levels of neovascularization at 12 weeks post-VML. (A-D) Representative images of vascular staining at 12 weeks post-VML. Images were taken at and around the defect site showing CD31^+^ cells (*red*, arrowhead) and αSMA^+^ structures (*green*, arrow) within the region of interest. (E-H) Magnified images of muscle defect areas in (E) native, (F) no repair, (G) CG scaffold, and (H) CG-PPy scaffold experimental groups. (I) CD31^+^ cell counts were significantly elevated in CG-PPy scaffold-treated muscles. (J) The number of αSMA^+^ structures was significantly increased in CG-PPy scaffold-treated muscles compared to all other experimental groups. Data presented as Mean +/-SD. ** *P* < 0.01, *** *P* < 0.001, **** *P* < 0.0001. *n* = 3 muscles per experimental group. Scale bars: 2 mm (A-D), 100 μm (E-H).

### 3.8 Assessment of muscle innervation

In addition to evaluating macrophage and vascular cell infiltration, IHC staining of neurofilament 200 (NF200) was used to probe for innervation at 12 weeks post-VML. Rapid skeletal muscle innervation following injury is necessary to facilitate functional recovery while also limiting prolonged denervation and muscle atrophy. NF200 structures localized toward the middle belly of the TA muscle and were generally observed in clusters surrounding muscle fibers (**Figure 8**). The presence of NF200 was significantly reduced in NR and CG-PPy scaffold-treated muscles compared to native tissues (NR: *P* = 0.025, PPy: *P* = 0.046), indicating limited innervation post-VML. In contrast, CG scaffold-treated tissues showed higher levels of NF200 structures that were similar to native muscle (CG: *P* = 0.467), corroborating the observed improved functional muscle contraction response to electrical stimulation.

**Figure 8:**
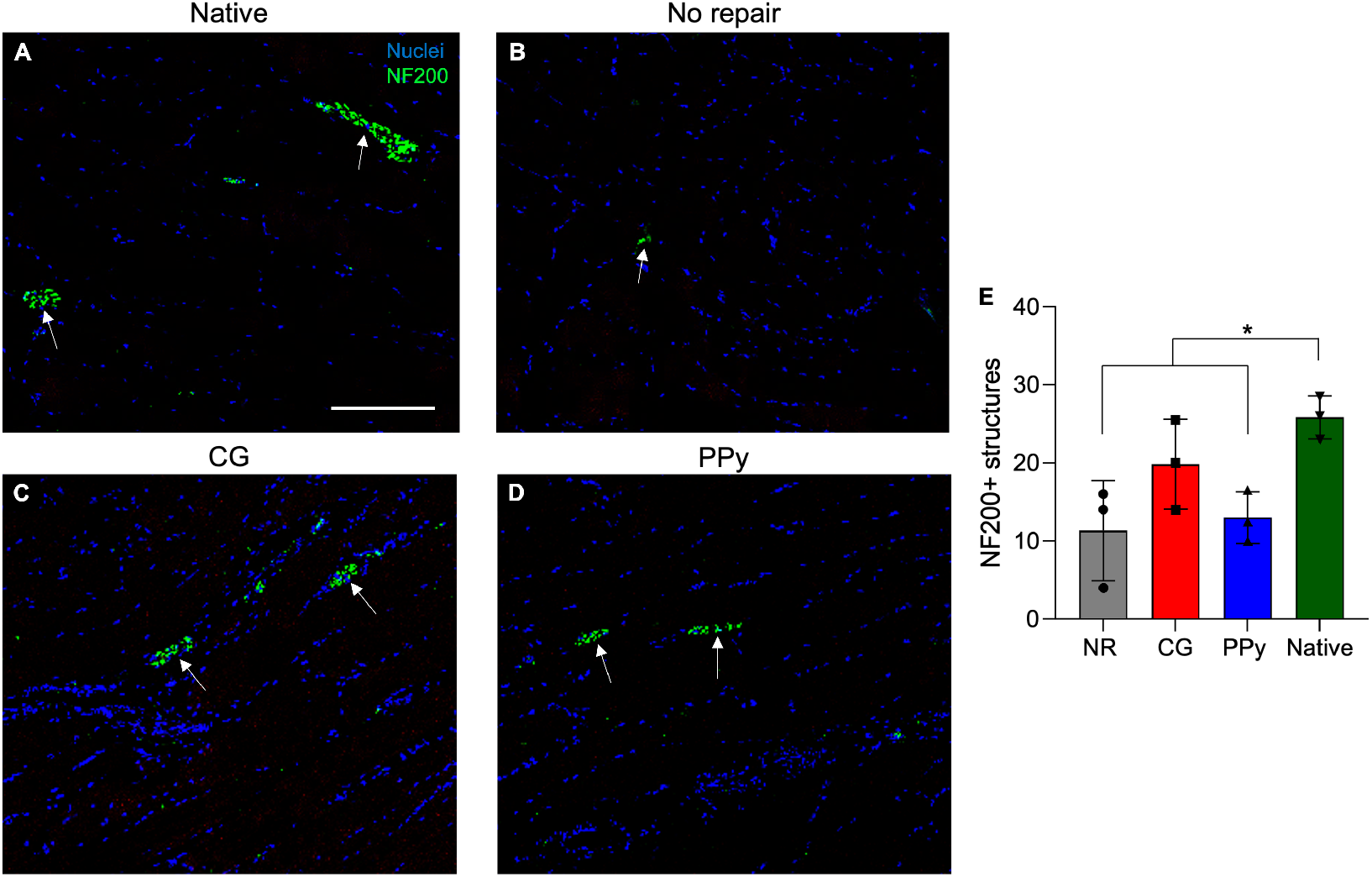
Muscle innervation is significantly reduced in non-treated muscles at 12 weeks post-VML. (A-D) Representative images of neurofilament staining at 12 weeks post-VML. NF200 structures (*green*, arrow) were quantified from images across longitudinal muscle sections. (E) Quantification of NF200 structures revealed that CG scaffold-treated tissues possess statistically similar levels of peripheral nerves compared to native muscle. Data presented as Mean +/-SD. * *P* < 0.05. *n* = 3 muscles per experimental group. Scale bar: 200 μm.

## 4. Discussion

The ideal therapeutic for VML injuries should facilitate restoration of muscle volume, complete with vasculature and innervation, and recovery of tissue function to preinjury levels. While recent advancements in biomaterial design and surgical techniques have shown promise for restoring muscle function in pre-clinical animal models, they remain limited in their ability to facilitate consistent clinical outcomes. As a result, a rigorous analysis of how material biophysical properties influence the body’s endogenous wound healing response is required to inform subsequent biomaterial design. Previous work has highlighted that material anisotropy and bioelectrical cues are important regulators of myoblast proliferation and maturation *in vitro*^38,39^, but it remains unclear how important these factors are in supporting endogenous cell-mediated repair *in vivo*. To address this knowledge gap, we explored how collagen scaffold anisotropy and conductivity impacted skeletal muscle wound healing in a biologically-relevant VML injury model.

We chose collagen as our material backbone due to its abundance as a component of the skeletal muscle ECM and previous use as a biomaterial platform for muscle tissue engineering^22,40,41^. Additionally, collagen-glycosaminoglycan (CG) scaffolds have been widely used across a range of different tissue engineering applications and are clinically approved for the treatment of skin and peripheral nerve defects^42^. To mimic the high level of organization observed in skeletal muscle, we employed a directional lyophilization technique using a thermally mismatched mold. The discontinuity in thermal properties within the mold allows for uniaxial heat transfer during freezing that results in alignment of ice crystals and anisotropy of the collagen struts^23^. All scaffolds were fabricated at a freezing temperature of -10°C and a cooling rate of 1°C/min, which has been shown to support aligned collagen struts, cell proliferation, and organized differentiation of myoblasts^18,23,43,44^. It is also important to note that naturally-derived biomaterial systems are typically limited by their suboptimal mechanical properties and are degraded rapidly by matrix metalloproteinases (MMPs). As a result, we dehydrothermally and chemically crosslinked the collagen and chondroitin sulfate within the scaffolds to increase resistance to enzymatic degradation and improve material mechanics^45^.

In an effort to simulate skeletal muscle’s inherit electrically responsive nature, we aimed to include a conductive moiety within our aligned CG scaffolds. The use of conductive polymers has been explored across a range of electrically-excitable cell types and has been shown to facilitate improved cell adhesion, proliferation^46,47^, and differentiation^48,49^. The most commonly used conductive polymers in biomaterial systems include polypyrrole (PPy)^50^, polyaniline (PANI)^51^, and poly(3,4-ethylenedioxythiophene) (PEDOT)^52^. We have previously shown that PPy in particular can be easily synthesized and incorporated into CG scaffolds, resulting in increased electrical conductivity^53–55^. Additionally, we previously showed that CG-PPy scaffolds supported myoblast growth, organization, and differentiation *in vitro*^18^. Previous work has also shown that PPy films result in minimal tissue toxicity and immune tissue response when implanted in a variety of animal models^56–58^. As a result, a logical extension of our prior work was to evaluate how CG scaffolds (with or without conductive PPy) support *in vivo* skeletal muscle wound healing.

Numerous different *in vivo* VML injury models have been evaluated in the literature with the majority focusing on repair of rodent limb muscles^31,59–61^. These models are beneficial because they provide a reasonable correlate to common civilian extremity trauma and combat-related injuries^62^. In this study, biomaterial efficacy was evaluated in a tibialis anterior (TA) model of VML. CG scaffolds could be easily manipulated and cut to the dimensions of the defect at the time of surgery to ensure complete defect coverage (**Figure 1**). Additionally, the shape of the scaffolds is only restricted by the geometry of the freeze-drying mold, and as a result scaffolds can be fabricated to treat a variety of complex and irregular injuries. Our results showed that both non-conductive CG scaffolds and conductive CG-PPy scaffolds facilitated improved functional muscle recovery as measured by *in vivo* maximum force generation compared to non-treated muscle (NR) at 12 weeks post-VML (**Figure 2**). Moreover, this trend was persistent when muscle force was normalized to baseline values, indicating functional recovery was not related to animal growth. It is important to note that synergistic muscles, extensor digitorum longus (EDL) and extensor hallucis longus (EHL), were surgically ablated to avoid compensatory hypertrophy following injury. Removal of these muscles results in an approximate 20% reduction in torque generation in the anterior compartment and as such, comparison of normalized torque post-injury would be theoretically limited to ∼ 80 N mm kg^-1^ (104.5 N mm kg^-1^ average at baseline). While maximal isometric contraction is an important metric to assess functional muscle recovery, restoration of submaximal muscle force is also an important clinical outcome to enable precise control of motor function. Muscle force remained statistically similar at lower stimulation frequencies (10-50 Hz) regardless of treatment type, suggesting that biomaterial design can be further optimized to improve functional outcomes. However, scaffold-treated groups produced superior muscle force at moderate frequencies (100-150 Hz) indicating collagen scaffolds supported improved muscle function and control. Interestingly, the non-conductive scaffolds displayed accelerated functional recovery compared to the NR animals as early as week 8. While this phenomenon invites further investigation, this could potentially be due to more rapid muscle cell infiltration and organization that is somewhat inhibited by residual PPy in the conductive scaffold group.

To elucidate the differences in cellular response to therapeutic intervention, experimental and contralateral control TA muscles were surgically explanted, and flash frozen for histological and immunohistochemical analysis. Unfortunately, freeze artifacts were observed in some muscle sections as circular gaps within muscle fibers, and future work will look to improve freezing methods. Hematoxylin and eosin (H&E) staining revealed a thin layer of regenerating muscle at the material-tissue interface, characterized by small, disorganized muscle fibers with centrally located nuclei (**Figure 4**), a finding that was not observed in non-treated tissues. These findings corroborate our functional testing data and suggest that both scaffold groups facilitated superior myogenesis that ultimately leads to improved muscle function. CG-PPy scaffold-treated muscles also showed residual conductive particles at the location of implantation. We have previously determined that PPy particles are approximately 500 nm in diameter and thus are not likely to be phagocytosed by cells. Our findings also reflect previously published work where an electrically responsive PPy-chitosan hydrogel was injected into the peri-infarct region of a rat heart 1 week after myocardial infarction^63^. In this study, the researchers observed that the PPy-chitosan hydrogel was still visible in the peri-infarct space and embedded in the repairing cardiac tissue of Masson trichrome-stained sections 8 weeks after injection. However, despite persistence of the conductive moiety at the injury site, PPy-chitosan gels facilitated increased transverse activation velocity and load-dependent ejection fraction fractional shortening in comparison with saline or chitosan injections^63^. Together these findings suggest that while PPy particles may remain at the site of injury 12 weeks post-injury, CG-PPy scaffold implantation may still support improved muscle function. Future work may look to improve cellular breakdown of conductive polymers by incorporating enzymatically degradable motifs, although such modification would likely decrease electrical properties.

It is important to note that despite evidence of myogenesis in scaffold-treated groups, a significant amount of fibrotic tissue remained, and all experimental tissues had significantly reduced muscle volume compared to native TA muscles (**Figure 3**). Previous work by our group and others have described the phenomenon of functional fibrosis in which some improvement in muscle function is observed, but very little *de novo* myogenesis occurs^64–66^. In this paradigm, the therapeutic mainly functions by providing passive mechanical force transmission along the remaining intact muscle approaching the theoretical maximal isometric contraction. In an effort to limit the development of functional fibrosis, the inclusion of myogenic cells within the regenerative therapeutic is likely necessary to restore muscle volume and function^31,64,67^. A meta-analysis exploring regenerative therapies following VML supports this hypothesis, stating that currently the most effective treatment for functional repair of skeletal muscle combines engineered biomaterials and seeded myogenic cells^67^. These findings are further supported by our group where a decellularized porcine bladder extracellular matrix (BAM) resulted in minimal functional muscle recovery and extensive fibrosis when implanted into a VML injury^64^. Conversely, when the BAM was reseeded with myogenic cells, substantial *de novo* myogenesis was observed, leading to superior functional recovery in numerous animal models^25,31,59^.

Although previous work highlights the importance of a cellular component to facilitate robust muscle regeneration, we chose to deploy the scaffolds acellularly due to the reduced regulatory challenges associated with these materials. Moreover, acellular CG scaffolds produced via freeze-drying are currently clinically approved for the treatment of skin and peripheral nerve injuries^21,68^. While our scaffold system did not appear to facilitate extensive myogenesis, scaffold-treated muscles showed evidence of regenerating myofibers and superior muscle force at sub-maximal electrical stimulation that was not observed in NR tissues. We believe that these enhanced regenerative outcomes are due to improved cell infiltration and remodeling at the site of VML. Freeze-dried CG scaffolds possess an open, interconnected pore microstructure (pore diameter ∼ 150 μm) that allows rapid cellular infiltration while facilitating nutrient and waste transport^18^. In contrast, traditional hydrogel systems contain nanometer-scale mesh sizes that are orders of magnitude smaller^23,68,69^ and inhibit these processes. In this initial study, cell and tissue organization was only characterized at 12 weeks post-injury, when scaffolds appeared to have fully degraded. Future work should assess cell infiltration into scaffolds at early time points post-injury to better evaluate endogenous repair. Despite this limitation, our findings highlight the use of CG scaffolds as potential therapeutics for VML injuries (**Table 1**).

**Table 1:**
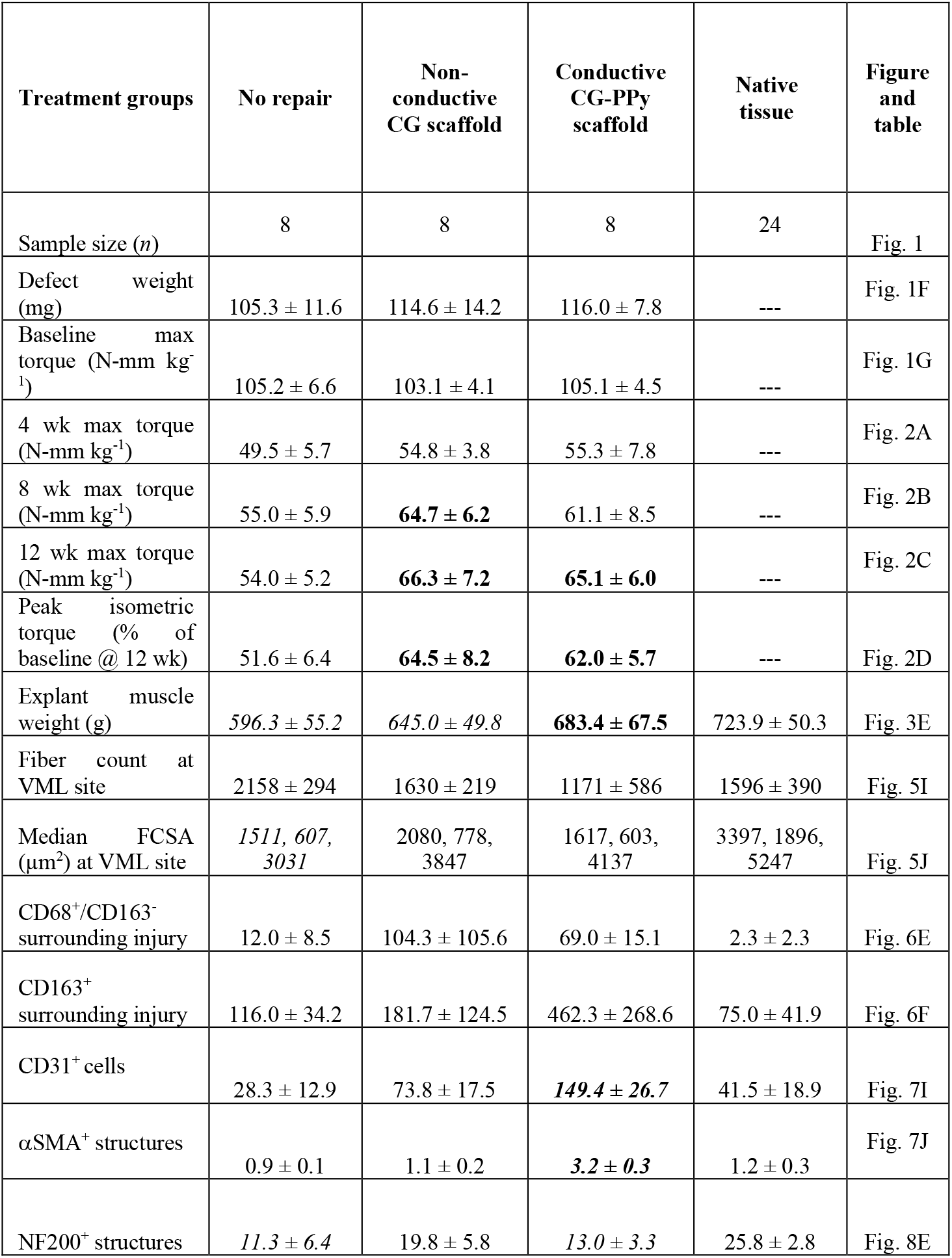
Summary of experimental findings across treatment groups. Values are presented as mean ± standard deviation except for median minimum FCSA, which is reported as the median, 1^st^, and 3^rd^ quartiles. Values denoted with the bolded lettering are significantly different (*P* < 0.05) from the no repair group, while values with italic lettering are significantly different (*P* < 0.05) from the native tissue contralateral control muscles. FCSA, fiber cross sectional area; NF200, Neurofilament 200; ---, analysis not applicable to native tissue contralateral control.

To further characterize the extent of skeletal muscle repair, SMASH analysis was used to quantify total myofiber number and fiber cross sectional area (FCSA) (**Figure S1**). Given the size of severity of the VML defect created there is minimal myogenesis in the non-repaired muscles, so the myofiber count for this experimental group is likely overestimated and more representative of fibers close to VML wound margin as opposed to newly-regenerated fibers. Analysis of myofiber counts showed that the number of muscle fibers was marginally reduced compared to native muscle tissues, indicating suboptimal recovery of muscle volume. This result further supports the claim that a cellular component is likely necessary to augment endogenous repair mechanisms to recover adequate muscle mass and function. Additional characterization of muscle FCSA at the VML injury revealed that median fiber size was significantly reduced in NR muscles compared native tissue (**Figure 5**). In contrast, median myofiber FCSA within scaffold-treated tissues was not statistically different from uninjured tissues, indicating that scaffolds facilitated superior muscle recovery. These findings support our functional data suggesting that the improvements observed in muscle torque production are related to improved cellular repair and wound healing, as myogenesis was not present in non-treated muscles. Despite improvements in median muscle FCSA, scaffold-treated tissues still displayed a leftward shift toward smaller muscle fibers compared to uninjured muscle. These findings suggests that while collagen scaffolds may support improved muscle function, they remain limited in their ability to facilitate extensive myogenesis. As a result, future work should look to improve repair of muscle volume using CG scaffolds by inclusion of myogenic and supporting cells or other bioactive moieties to further augment endogenous repair mechanisms.

VML injuries are characterized by a persistent inflammatory response which limits myogenic cell expansion and instead facilitates myofibroblast activation, ECM deposition, and fibrosis. While many inflammatory cells are involved in skeletal muscle repair, macrophages have been highlighted as important mediators of the wound healing cascade that can either serve as key constituents of pro-regenerative microenvironments or perpetuate chronic inflammation^35,36^. During normal wound healing, a carefully controlled shift in macrophage polarization from a classically-activated M1 ‘pro-inflammatory’ phenotype toward a predominately alternatively-activated M2 ‘pro-regenerative’ phenotype allows for robust myoblast expansion followed by differentiation and maturation^35–37^. However, during VML injuries macrophages are persistently activated, thereby limiting endogenous repair mechanisms and instead promoting activation of cells from fibrogenic lineages^3,70^. By evaluating macrophage infiltration and polarization within the VML injury area we aimed to assess the scaffold’s ability to support a pro-regenerative microenvironment (**Figure 6**). The number of M1 (CD68^+^/CD163^-^) and M2 (CD163^+^) macrophages remained elevated in CG-treated muscles (M1: *P* = 0.169; M2: *P* = 0.825) compared to native muscle, suggesting a persistent inflammatory response. While not statistically significant, CG-PPy scaffold-treated muscles also showed an increase in both M1 and M2 macrophages, indicating continual remodeling (M1: *P* = 0.467; M2: *P* = 0.054). Interestingly, non-treated muscles showed macrophage staining profiles most similar to native tissues. Typically, during wound healing macrophages return to baseline values at approximately 2 weeks post-injury to allow for the repair and remodeling phases. As a result, we hypothesize that the wound healing response reached completion prior to 12 weeks in NR tissues, leading to a reduction in number of macrophages and the presence of necrotic muscle. Future work would benefit from exploring earlier time points post-injury and assessment of additional cell types involved in aberrant wound healing, such as fibroadipogenic progenitor cells, to gain a more complete picture of the repair process.

After investigating macrophage localization, we next quantified the number of endothelial cells (CD31^+^) and pericytes (αSMA^+^ cells co-localized with CD31^+^ endothelial cells) as a proxy of vascularization (**Figure 7**). Given the high metabolic needs of skeletal muscle, angiogenesis and re-vascularization following VML are integral to functional recovery^71^. Significantly higher numbers of CD31^+^ cells and αSMA^+^ structures were observed in the CG-PPy scaffold-treated muscles compared to the other experimental groups. While other work has shown that PPy materials can enhance vascularization^72,73^, our observed results may be due to the overall higher level of cellularity in the PPy group. Future work could further improve vascularization by tuning scaffold properties such as glycosaminoglycan content, where previous work incorporating heparin significantly improved angiogenic outcomes^29,74,75^.

While the restoration of muscle volume is essential for improved regenerative outcomes, innervation of repaired tissue is also required for proper function. Electrically-responsive polymers have previously been shown to facilitate increased neural stem cell proliferation and differentiation *in vitro* compared to non-conductive substrates^58,76^. Therefore, we aimed to assess if scaffolds allowed superior skeletal muscle innervation *in vivo* compared to other experimental groups. Myofibers are innervated at neuromuscular junctions (NMJs) that are often found along the sarcolemma toward the middle of the muscle belly to allow for effective propagation of action potentials^77^. As a result, TA muscles were cut in half, sectioned longitudinally, and stained for neurofilament 200 (NF200) to visualize neurons (**Figure 8**). Only non-conductive CG scaffolds contained statistically similar levels of NF200^+^ structures compared to native muscle. In contrast, no repair and conductive CG-PPy scaffold-treated tissues possessed significantly reduced numbers of NF200^+^ structures. These findings corroborate our functional testing data in which only CG scaffold-treated muscles produced increased isometric torque at sub-maximal stimulation frequencies of 60 and 80 Hz. We hypothesize that the lack of innervation in CG-PPy scaffold-treated muscle may be due to reduced electrical dopant stability under physiological conditions. Previous work has shown that substrate conductivity can decrease substantially over time reducing the electrical properties of the scaffold^78^. The findings further indicate that while conductive PPy may facilitate improved cell development *in vitro*, further material optimization is required for *in vivo* efficacy.

## 5. Conclusion

This study aimed to improve the treatment of VML injuries by designing a biomaterial system that mimics biophysical features critical to healthy function, including three-dimensional (3D) anisotropy and electrical excitability. In the present study, we assess the efficacy of a 3D aligned collagen scaffold platform to repair a biologically-relevant TA model of VML injury. Non-conductive CG and electrically conductive CG-PPy scaffolds were prepared by directional lyophilization and surgically implanted into VML defects. Both scaffold-treated groups supported increased functional muscle recovery at 12 weeks post-injury compared to non-treated muscles as measured by *in vivo* electrical stimulation of the peroneal nerve. Subsequent histological analysis of the material-tissue interface showed regenerating muscle fibers in scaffold-treated muscles, indicated by small myofibers with centrally located nuclei, that was not observed in non-repaired tissues. Further analysis of myofiber size showed that non-treated tissues possessed significantly reduced median FCSA compared to native muscles, indicating impairment of muscle regeneration. Immunohistochemical analyses revealed that scaffold-treated muscles possessed higher numbers of macrophages and conductive scaffold-treated muscles had higher numbers of vascular cells, suggesting a persistent wound healing response that was not observed in non-treated tissues. Finally, non-conductive scaffolds alone possessed statistically similar levels of neurofilament staining compared to native tissue, further corroborating the improved isometric contraction observed. While the addition of PPy largely did not appear to result in improved functional outcomes compared to non-conductive CG scaffolds, our findings illustrate that aligned collagen scaffolds can facilitate improved functional recovery and endogenous muscle repair following VML injury. Future work should look to optimize the synthesis of CG scaffold composites to further augment skeletal muscle regeneration.

## Supporting Information

Additional data on measurements of muscle fiber cross-sectional area and minimum Feret diameter can be found in the Supporting Information.

## Acknowledgments

The authors would like to acknowledge the University of Virginia’s Histology Core facility for help with histological sectioning, Gregg Gardner for his help quantifying muscle fiber number and cross-sectional area, and Michael Rariden for helpful advice regarding muscle fiber characterization data. This work was supported by seed funding from the University of Virginia’s Center for Advanced Biomanufacturing, the UVA Biotechnology Training Program (T32GM136615), and the NIH (R21AR075181 and R01AR078886).

The content is solely the responsibility of the authors and does not necessarily represent the official views of the National Institutes of Health.

## Notes

### Competing Interest Statement

The authors have declared no competing interest.

