## Supporting Information for "Freeze-dried porous collagen scaffolds for the repair of volumetric muscle loss injuries"

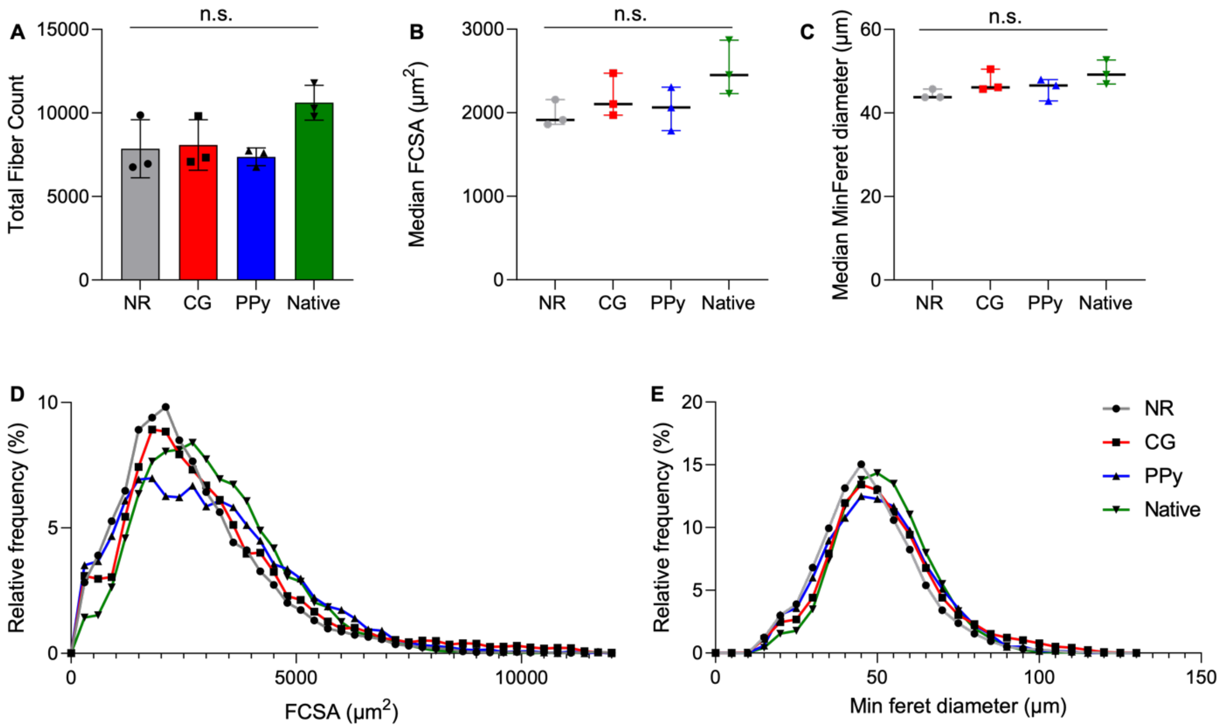


**Figure S1: Total myofiber number and cross-sectional area is reduced regardless of treatment type.** (A) Total myofiber count was not statistically different from native muscle across experimental groups, although the number of muscle fibers was reduced. (B) Total median fiber cross-sectional area (FCSA) and (C) median minimum Feret diameter were statistically similar across all experimental groups. (D) FCSA and (E) minimum Feret diameter relative frequency curves show similar distributions of myofiber size regardless of treatment type. Data presented as Mean +/- SD while panel (B) and (C) data are presented as the median (line: median) with interquartile range (whiskers). n.s.: no statistically significant differences. *n* = 3 muscles per experimental group.

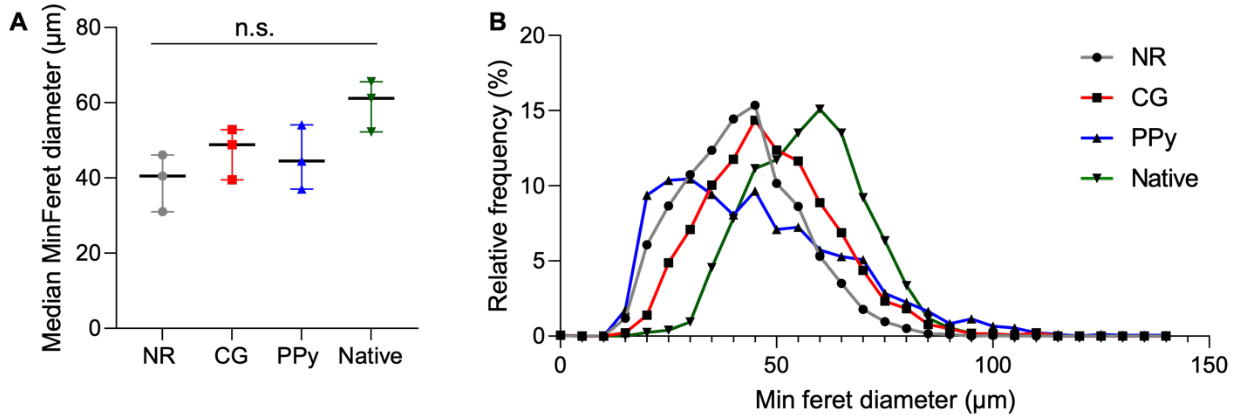


**Figure S2: Myofiber minimum Feret diameter is reduced in non-treated muscle tissues at the site of VML injury.** (A) Minimum Feret diameter was reduced in NR muscles, although these results were not statistically significant. (B) Minimum Feret diameter relative frequency curves show a similar leftward shift toward smaller muscle fibers across experimental tissues compared to native muscles. Panel (A) data are presented as the median (line: median) with interquartile range (whiskers). n.s.: no statistically significant differences. $n$ = 3 muscles per experimental group.
